# Codon optimisation for maximising gene expression in multiple species and microbial consortia

**DOI:** 10.1101/2020.06.30.177766

**Authors:** David J. Skelton, Lucy E. Eland, Martin Sim, Michael A. White, Russell J. Davenport, Anil Wipat

## Abstract

**Motivation:** Codon optimisation, the process of adapting the codon composition of a coding sequence, is often used in synthetic biology to increase expression of a heterologous protein. Recently, a number of synthetic biology approaches that allow synthetic constructs to be deployed in multiple organisms have been published. However, so far, design tools for codon optimisation have not been updated to reflect these new approaches.

**Approach:** We designed an evolutionary algorithm (EA) to design coding sequences (CDSs) that encode a target protein for one or more target organisms, based on the Chimera average repetitive substring (ARS) metric — a correlate of gene expression. A parameter scan was then used to find optimal parameter sets. Using the optimal parameter sets, three heterologous proteins were repeatedly optimised *Bacillus subtilis* 168 and *Escherichia coli* MG1655. The ARS scores of the resulting sequences were compared to the ARS scores of coding sequences that had been optimised for each organism individually (using Chimera Map).

**Results:** We demonstrate that an EA is a valid approach to optimising a coding sequence for multiple organisms at once; both crossover and mutation operators were shown to be necessary for the best performance. In some scenarios, the EA generated CDSs that had higher ARS scores than CDSs optimised for the individual organisms, suggesting that the EA exploits the CDS design space in a way that Chimera Map does not.

**Availability and implementation:** The implementation of the EA, with instructions, is available on GitHub: https://github.com/intbio-ncl/chimera_evolve.

## 1. Introduction

Codon optimisation is the process of adapting the codon composition of a coding sequence towards a given end, normally increasing the expression of a heterologous protein. The relationship between a coding sequence (CDS) and the expression of the encoded protein is complex, and not fully understood [1].

The earliest codon optimisation methods leveraged a one amino acid-one codon approach to codon optimisation, in which every amino acid is assumed to be encoded by a single, optimal codon. Methods using this approach include optimising for the codon adaptation index (CAI) [2], and relative codon adaptation (RCA) [3] measures. In CAI, each codon in a CDS is given a score, *w*_*i*_, as shown in equation 1, where *f*_*i*_ is the frequency of the codon in a reference set of highly expressed sequences, and *max*(*f*_*j*_) is the frequency of the most frequent synonymous codon in the same set. The CAI of the CDS is then the geometric mean of all the codon scores.

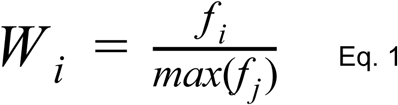

A potential issue with one amino acid-one codon approaches is that, being based on single-codon distributions, they often do not capture the emergent properties of the resulting macromolecules [4]. For example, there have been suggestions that protein folding is related to the rate of tRNA binding [5] and mRNA secondary structure, both of which are affected by the codon composition of the CDS [6]. Consequently, newer methods utilise other approaches to optimise CDSs.

Another method, Chimera Map, constructs coding sequences using codon “blocks” from native coding sequences, aiming to optimise a metric known as the Chimera Average Repetitive Substring (ARS) score [1]. The ARS of a CDS is calculated as follows: each codon in the coding sequence (*S*) is given a score 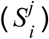, which is the length of the longest substring in *S*, starting at position *i*, that also appears in a CDS in a reference set. The ARS score is the sum of these scores divided by the length of *S* (Eq 2).

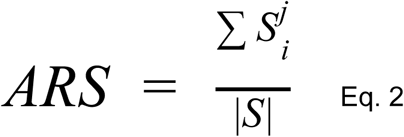

In their 2015 paper, Zur and coworkers demonstrated that the Chimera ARS score correlates better than CAI with the translation rates of heterologous sequences in *E. coli*. Crucially, the ARS score is calculated from a reference set comprising all CDSs in a target organism’s genome, rather than just highly-expressed CDSs. This is advantageous because it enables codon optimization for incomplete genomes and metagenomes as long as some CDSs can be called in the assembly, regardless of what they encode.

As the goals of synthetic biology become more ambitious, the methods employed for codon optimisation need to be adapted to meet a new set of requirements. Recently, projects to design genetic systems that are deployed in multiple species or multicellular systems have become more common. An example of this approach is the use of integrative conjugative elements to transfer DNA amongst microorganisms in consortia [7]. Other projects, such as the Portabolomics project at Newcastle University, aim to construct portable genetic systems that can operate in different hosts without the need for refactoring. In terms of codon optimisation, these two use-cases potentially represent different optimisation goals: one designing a construct that optimises the behaviour of a system, and the other designing a construct that will work well in a selection of target hosts.

Codon optimization algorithms, therefore, must be updated to ensure the maximal expression of synthetic CDSs in multiple target organisms. Unlike single organism optimisations, optimising a CDS for multiple organisms is a multi-objective optimisation problem, and can rapidly become intractable to deterministic approaches. Heuristics such as evolutionary algorithms have proven useful to tackle this kind of challenge in previous studies.

In this study, we demonstrate that an evolutionary algorithm (EA) is a feasible approach for the simultaneous optimisation of a single CDS for two species, using the Chimera ARS metric. Additionally, we demonstrate that the Chimera ARS scores for coding sequences can be optimised beyond what is achievable using Chimera Map.

## 2. Methods

### 2.1 Implementation of the evolutionary algorithm

In this section we describe an EA for optimising a single target protein sequence for one or more target organisms. A target protein is a single amino acid sequence of any length, with or without a stop codon. A target organism is a specific organism in which the CDS being designed by the EA is intended to be implemented.

The EA was implemented in Rust, and takes two inputs: (A) a target protein; and (B) one or more files of coding sequences, each of which contains the native CDSs from a single target organism. First, substrings from each target organism that could be used to encode part of the target protein are identified and filtered to remove redundancy. The codon strings are then converted into a compressed representation, in which each codon is replaced with a single character, and stored in a suffix table for fast searching.

The algorithm proceeds based on four parameters: *C*, the number of crossover events per generation; *M*, the number of mutation events per generation; *G*, the number of generations; and *S*, the size of the population at which to stop binary tournament selection. The first generation of the algorithm starts by generating *C* × 2 + *M* random candidate solutions encoding the target protein, which are then scored (Section 2.1.1). The number of candidates is then reduced through binary tournament selection: while the population size is greater than *S*, two members of the population are selected through random uniform selection, and the member with the lowest score is removed from the population. Binary tournament selection ensures that the candidate with the highest score in a pair remains in the population, but also gives lower scoring candidates an opportunity to persist, allowing the algorithm opportunities to explore new regions of sequence space.

For the remaining (*G* − 1) generations, each generation begins with *C* crossover events. Two random members of the population, *A* and *B*, are selected, and two random positions across the length of the candidate CDS, *P* and *Q* are selected, both using uniform selection. Two new candidates are generated by exchanging the region selected in the two candidates, `A[→ P] + B[P → Q] + A[Q →]` and `B[ → P] + A[P → Q] + B[Q →]` (Figure 2).

**Figure 1:**
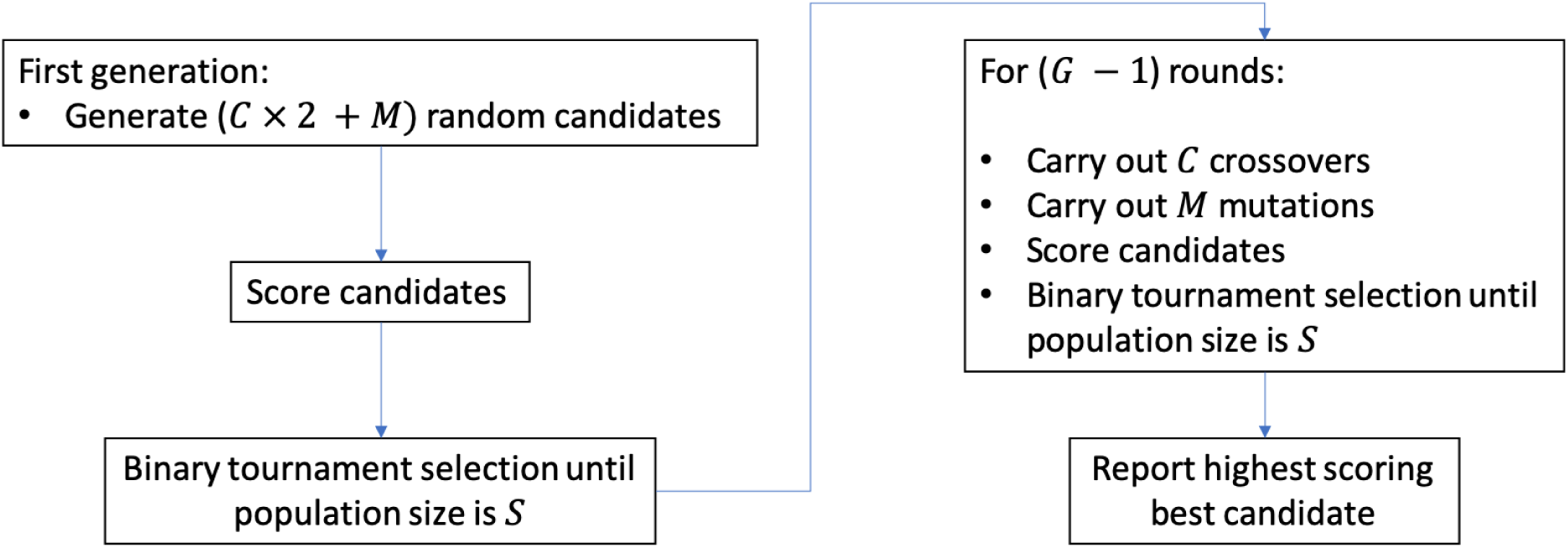
Procedure of the EA. Each generation consists of three stages: candidate generation, scoring, and tournament selection. In the first generation, all candidates are randomly created. In all remaining generations, candidates are generated using crossover and mutation operators.

**Figure 2:**
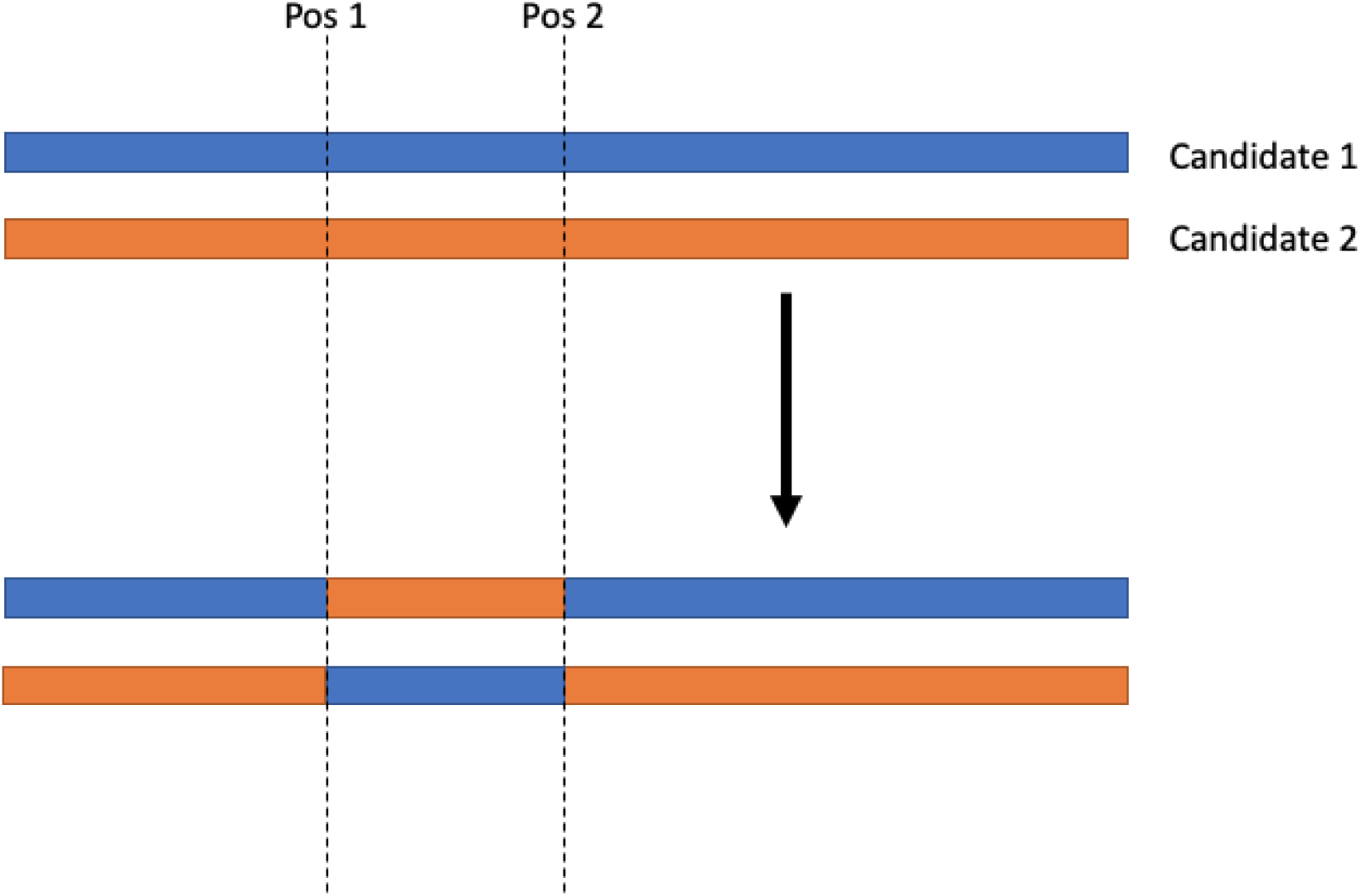
Example of a crossover event, demonstrating how two new candidates are generated from two original candidates and two positions for the crossover to occur.

Following crossover, *M* mutation events occur. In a mutation event, a random member of the population is selected, and between one and five codons are selected to be mutated. A codon can only be selected to be modified if an alternative codon encoding the same amino acid exists; in translation table 11, the codons ATG (methionine) and TGG (tryptophan) cannot be selected for mutation. For start codons, the current implementation of the EA only selects ATG. The selected codons are then exchanged for another codon encoding the same amino acid, generating a new candidate.

The generation is then completed by scoring unscored candidates (i.e., those that did not persist from the previous generation), and binary tournament selection is carried out as in generation one.

#### 2.1.1 Scoring

The algorithm was run in two modes: *minimum* or *weighted*, both of which use the Chimera ARS score.

In minimum mode, the Chimera ARS of a candidate CDS was calculated for each target organism, and the lowest of these was taken to be the fitness score of the candidate. In weighted mode, each target organism was given a user-specified weight, and the fitness score of a candidate was the normalised sum of the weighted Chimera ARS scores.

### 2.2 Identification of an optimal parameter set

One hundred random protein sequences were obtained from SwissProt [8] using the UniProt programmatic endpoint. Each sequence was optimized for *Bacillus subtilis* 168 and *Escherichia coli* MG1655 using all combinations of (A) parameter set (Table 1), (B) algorithm mode [*minimum, weighted*], and (C) generation start size in [100, 200]. In weighted mode, *B. subtilis* and *E. coli* were equally weighted. The parameter sets in Table 1 were chosen so that, in total, 500,000 candidate coding sequences would be generated:

**Table 1:** Parameter settings used to test the evolutionary algorithm. Each parameter setting was run once per algorithm mode (minimum and weighted) and generation start size (100 and 200).

| Number of mutation events / generation | Number of crossover events / generation | Number of generations |
| --- | --- | --- |
| 0 | 250 | 1,000 |
| 100 | 200 | 1,000 |
| 200 | 150 | 1,000 |
| 300 | 100 | 1,000 |
| 400 | 50 | 1,000 |
| 500 | 0 | 1,000 |
| 0 | 500 | 500 |
| 200 | 400 | 500 |
| 400 | 300 | 500 |
| 600 | 200 | 500 |
| 800 | 100 | 500 |
| 1,000 | 0 | 500 |

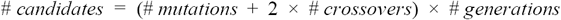

To facilitate comparisons, parameter sets were chosen so that 500,000 candidates would be generated across a single run of the EA.

For each algorithm mode, the results of the 24 different settings were compared using the Wilcoxon signed-rank test (`scipy`). The best parameter sets were those that never had a statistically lower population mean rank than the results of another parameter setting.

### 2.3 Evaluation of optimisation consistency

For each algorithm mode, the best performing parameter sets identified in section 2.2 were used to optimise three proteins: green fluorescent protein from *Aequorea victoria* (P42212), luciferin 4-monooxygenase from *Luciola mingrelica* (Q26304), and CRISPR-associated endonuclease Cas9/Csn1 from *Streptococcus pyogenes* serotype M1 (Q99ZW2). Optimisations were repeated 10 times to obtain a distribution of scores for comparison.

## 3. Results

### 3.1 Identification of the best performing parameter sets

The optimal parameter set experiments took a total of 31:22:07 (HH:MM:SS) in weighted mode (median time per optimization was 25s, IQR 13.7s - 42.6s) and 28:41:37 in minimum mode (median time per optimization was 22.9s, IQR 13.1s - 39.2s). Experiments were carried out on a server with 32 threads (Intel Xeon Processor E5-2670) and 126 GB RAM. It should be noted that these resources far exceed what is required for a single optimisation (a standard desktop is sufficient), and were used to speed up parallelizable steps within the EA.

Figure 3 shows the ranked results of using 24 different parameter settings (plus random) to generate coding sequences for 100 proteins with the algorithm minimum mode. Three parameter settings were found to be best performing (table 2). Of note, it can be seen that parametrizations with zero mutations or zero crossover events were amongst the worst performing. In addition to having a lower rank, parameter sets with zero mutation operators, had a noticeably lower absolute score (Figure 4).

**Table 2:** Best performing parameter sets (by mode type). A parameter set was said to be best performing if a Wilcoxon signed rank test against the results of all other parameter settings never found a median of differences to be statistically significantly negative.

| Mutations | Crossovers | Generations | Generation start size |
| --- | --- | --- | --- |
| Minimum mode |  |  |  |
| 300 | 100 | 1,000 | 200 |
| 400 | 50 | 1,000 | 200 |
| 800 | 100 | 500 | 200 |
| Weighted mode |  |  |  |
| 300 | 100 | 1,000 | 200 |
| 400 | 50 | 1,000 | 100 |

**Figure 3:**
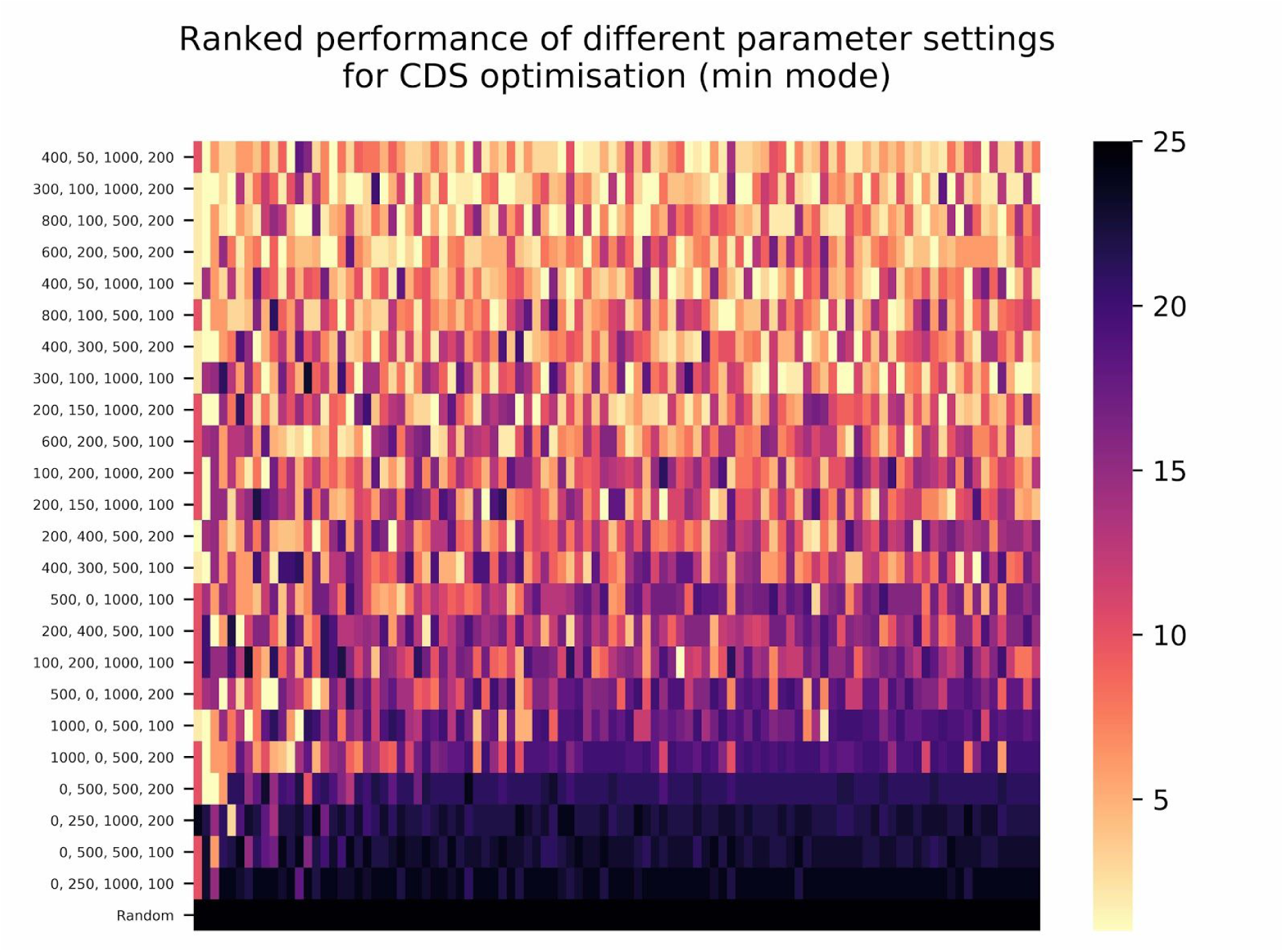
Ranked scores from running single optimisations in minimum mode (1 is best scoring, 25 is lowest scoring). Y axis plots the parameter settings as (# mutations, # crossovers, # generations, and # generation start size), and each column represents a different, randomly selected protein from UniProt.

**Figure 4:**
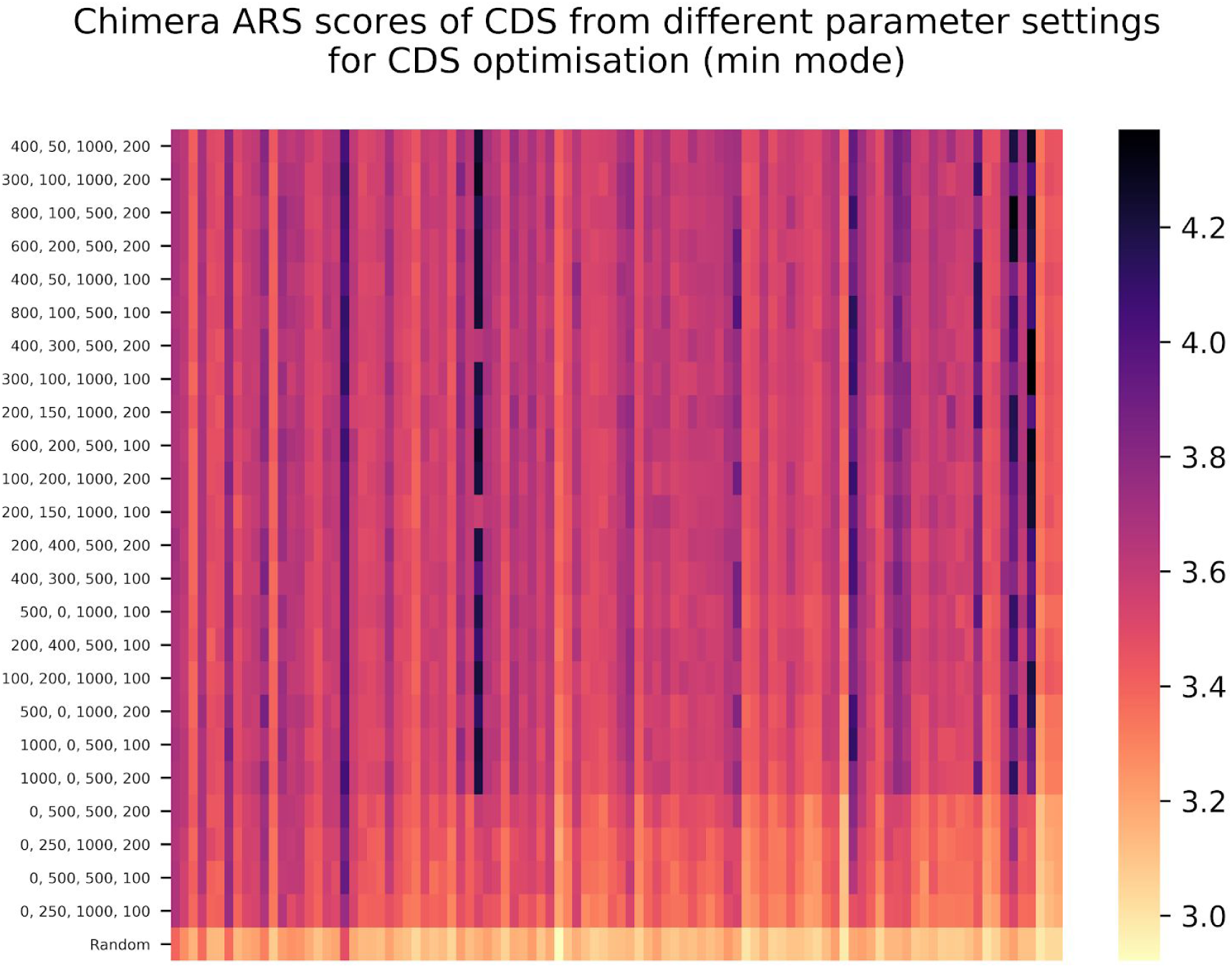
Raw Chimera ARS scores from running single optimisations in minimum mode. Y axis plots the parameter settings as (# mutations, # crossovers, # generations, and # generation start size), and each column represents a different, randomly selected protein from UniProt.

Figure 5 shows the results of the 24 parameter optimisation for 100 sequences in weighted mode. There were two sets of parameters that were found to be best performing from this experiment (Table 2), one of which was identical to a best-performing parameter set in minimum mode. Unlike minimum mode, there were some proteins that multiple parameter settings were able to optimize equally well (such as B7LUL6, the 50S ribosomal protein L7/L21 from *Escherichia fergusonii*).

**Figure 5:**
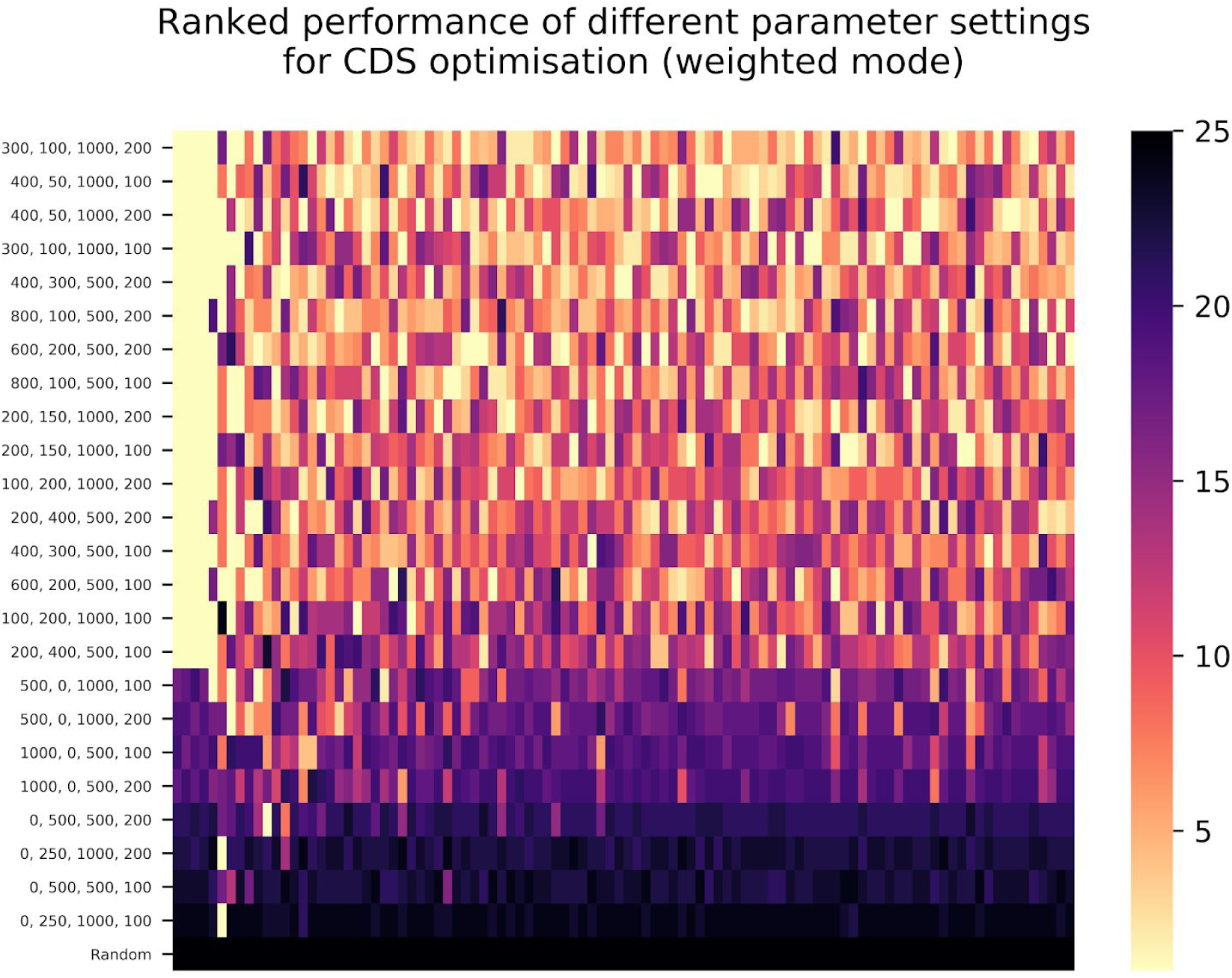
Ranked scores from running single optimisations in weighted mode (1 is best scoring, 25 is lowest scoring). Y axis plots the parameter settings as (# mutations, # crossovers, # generations, and # generation start size), and each column represents a different, randomly selected protein from UniProt. In this mode, there were often ties for best scoring construct (e.g., the first five columns).

The raw chimera ARS scores (Figure 6) demonstrate that the possible ARS score achievable is highly protein dependent. The two proteins with the highest achieved ARS scores were P39817, a Proton / glutamate-aspartate symporter from *B. subtilis* strain 168, and B7NV70, a CTP synthase from *E. coli* O7:K1. It is likely, therefore, that one of the organisms used for the experiment had close homologues to these proteins. Due to the non-linearity of the Chimera ARS score, the EA could optimise heavily for that organism with the homologue, consequently achieving a high weighted ARS score.

**Figure 6:**
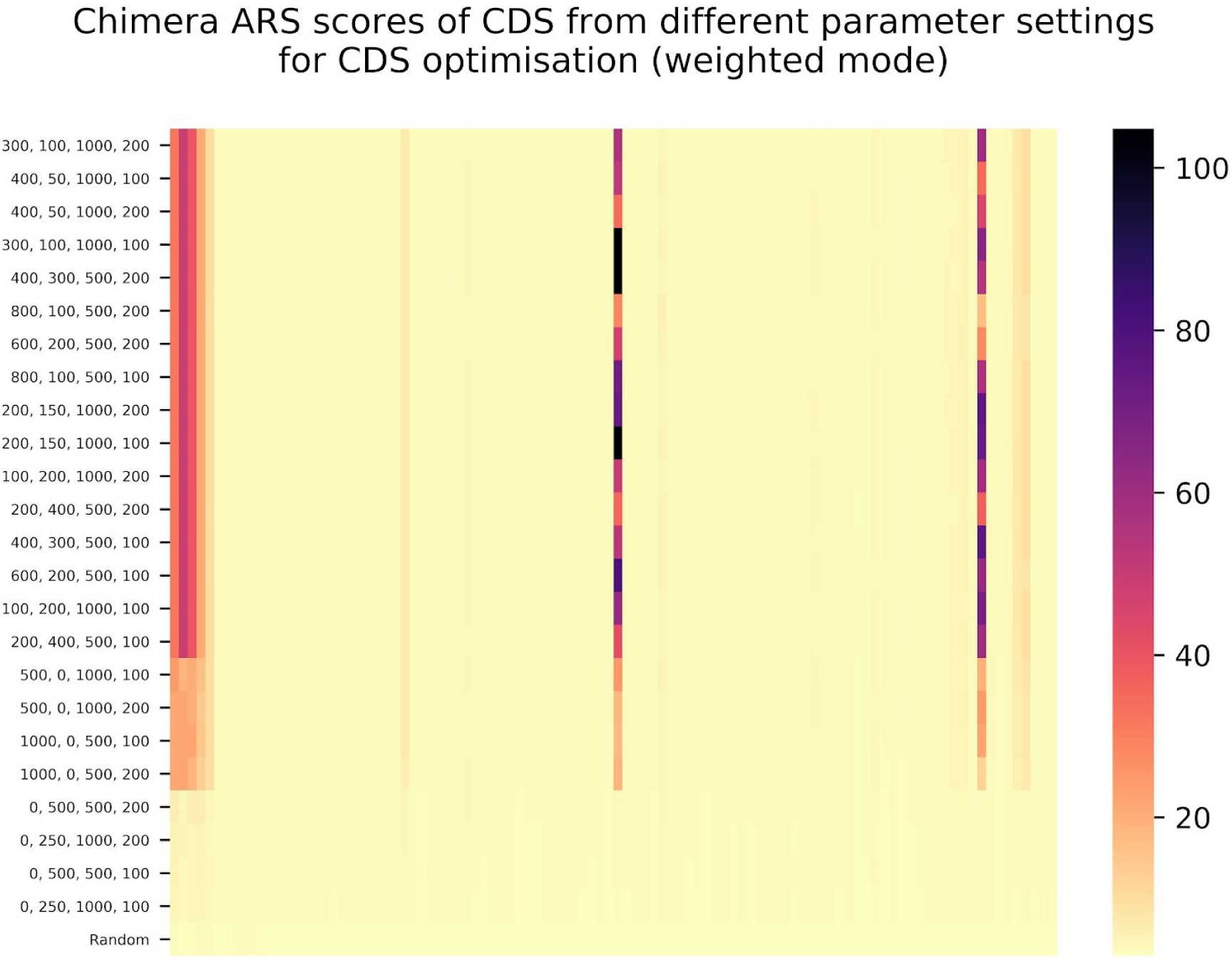
Raw Chimera ARS scores from running single optimisations in weighted mode. Y axis plots the parameter settings as (# mutations, # crossovers, # generations, and # generation start size), and each column represents a different, randomly selected protein from UniProt. More extreme scores were observed in this mode (note the scale on the colour bar).

### 3.2 Consistency results

The scores obtained for repeated optimisations are shown in Figure 7. In all cases, it can be seen that the results of the different parameter settings overlap, although the ranges of scores obtained do appear to differ by both parameter setting and protein. In some cases, the scores of the CDSs obtained exceeded scores obtained by designing constructs individually for each organism with Chimera UGEM. In the case of the highest-scoring construct for GFP (P42212) in weighted mode using 300 mutations, 100 crossovers, 1,000 generations and a generation start size of 200, the optimal construct scored 6.6303 (an average of the ARS score of 3.6176 for *B. subtilis* and 3.6429 for *E. coli* -- both higher than the ARS of constructs designed individually).

**Figure 7:**
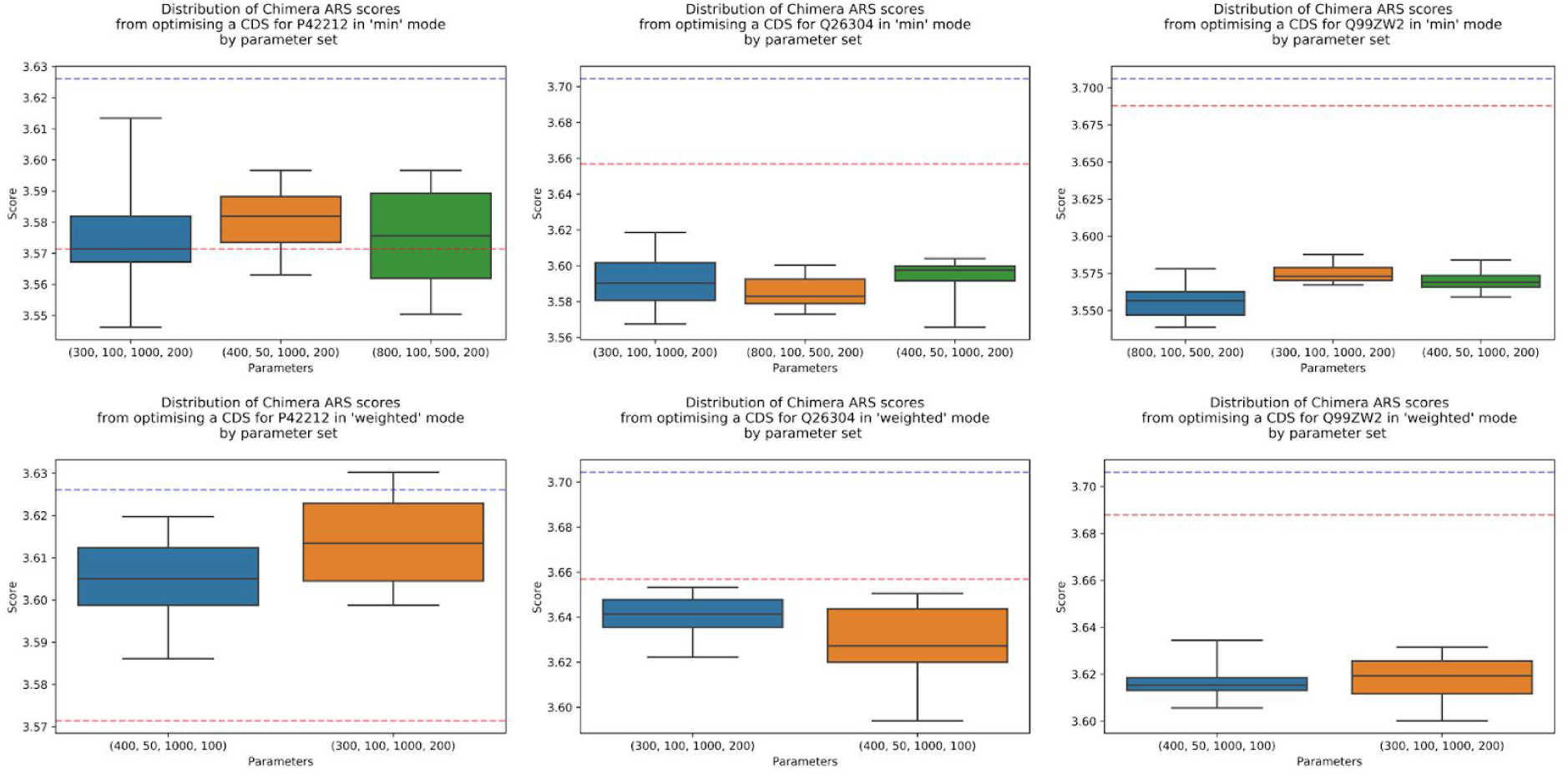
Scores achieved from running the evolutionary algorithm repeatedly to optimise a CDS encoding P42212, Q26304, and Q99ZW2 for *B. subtilis* and *E. coli* in minimum mode (above) and weighted mode (below). Parameter sets chosen were the optimal parameters obtained in section 3.1. Scores in the upper figure are the minimum ARS of the construct for either *E. coli* or *B. subtilis* (whichever was lower), whereas in the lower figure the score is the average. Blue line in the figures indicates the Chimera ARS score of the optimal construct designed by Chimera UGEM (window size = 0) and the red line indicates the same for *B. subtilis*. In the case of P42212, the scores obtained counterintuitively exceeded the optimal scores obtained for one or both of the organisms individually.

## 4. Discussion

This work, as far as the authors are aware, is the first instance of software being designed to optimise a single CDS for multiple organisms simultaneously. We presented an EA that can optimise coding sequences in two different modes -- demonstrating that the EA is able to consistently achieve scores similar to equivalent software aimed at single-species CDS optimisation, and in some cases out-perform this software.

The two modes implemented here were designed to address two separate anticipated use-cases. Minimum mode was designed to optimise a single CDS that expresses well in each of the organisms being optimised for. Thus, minimum mode prevents a CDS from being optimised for one organism at the expense of another. Conversely, weighted mode was designed to allow selective emphasis on certain species in a population, and is anticipated to be useful for optimising a sequence for microbial consortia (where composition may not be uniform). In weighted mode, we identified that designing a CDS encoding a homologous protein to a protein from target organisms leads to the algorithm optimising towards those variants (subject to the parameterisation). Whether this behaviour is desirable or not is unknown; it would be beneficial for future studies to include *in vivo* experiments to explore the relationship between (A) construct score(s), (B) weighting of the EA, (C) abundance of microorganisms in microbial consortia, and (D) expression characteristics of the microbial consortia.

These results also demonstrate that Chimera UGEM does not design the construct with the optimal Chimera ARS score -- if this were the case, then the horizontal line representing the ARS score of the UGEM designed GFP CDS for *B. subtilis* would be the ceiling for a construct designed in minimum mode. This likely reflects that the EA is able to explore solutions in a design space not considered by the Chimera Map implementation used by Chimera UGEM. In particular, the Chimera Map algorithm attempts to minimize the number of blocks used to construct a sequence, whereas we hypothesise that an EA also optimises the overlap space between blocks. Whether this is genuinely desirable behaviour or not is an open question and should be investigated further, with refinement of the Chimera ARS measure, if appropriate.

The results from the parameter set optimisations, while different between minimum mode and weighted mode, do share one parameter set in common: 300 mutations, 100 crossover events, 1,000 generations, and a generation start size of 200, suggesting that this parameter set could make an ideal default (regardless of mode). Crucially, none of the optimal parameter sets have zero crossover events or zero mutation events, suggesting that both of these operators contribute towards generating constructs with higher Chimera ARS scores. Further, the ratio of mutations to crossover events is similar in all of the optimal parameter sets (300:100, 400:50, and 800:100), with some dependence on the generation start size that appears to differ between modes.

While this work evaluates many of the parameter sets, further investigation of the evolutionary algorithm could yield a superior algorithm. For example, all of the randomisation in this study is random uniform selection, the maximum number of mutations occurring in a single mutation event was fixed at five, and selection of candidate CDSs between generations is fixed as binary tournament selection. While systematically exploring parameterisations and alternative protocols for these would have been ideal, further combinatorial explosion combined with an already expensive execution time made this impractical for the current study.

More recently, Diament *et al*. have developed a newer version of the Chimera Map algorithm, known as Chimera UGEM, that additionally takes into consideration where codon blocks occur within a coding sequence [9]. Thus, when building on our work, future work should consider how to incorporate the position-specific approaches put forward by Diament *et al*.

Finally, future research should be supplemented with *in vivo* experiments that would enable validation of the *in silico* results. Furthermore, *in vivo* experiments may guide the use of the EA presented here. For example, in weighted mode, there is still an opportunity to optimise further the weightings to maximise population-wide expression.

## Acknowledgements

The authors gratefully acknowledge funding from the Newcastle University Frontiers in Engineering Biology (NUFEB) project, EP/K039083/1, and the Portabolomics project, EP/N031962/1.

